# The Cell Cycle Browser: an interactive tool for visualizing, simulating, and perturbing cell cycle progression

**DOI:** 10.1101/234039

**Authors:** David Borland, Hong Yi, Gavin D. Grant, Kasia M. Kedziora, Hui Xiao Chao, Rachel A. Haggerty, Jayashree Kumar, Samuel C. Wolff, Jeanette G. Cook, Jeremy E. Purvis

**Affiliations:** Renaissance Computing Institute, University of North Carolina, Chapel Hill, 120 Mason Farm Road, Chapel Hill, NC 27599-7264; Department of Biochemistry and Biophysics, University of North Carolina, Chapel Hill, 120 Mason Farm Road, Chapel Hill, NC 27599-7264; Department of Genetics, University of North Carolina, Chapel Hill, 120 Mason Farm Road, Chapel Hill, NC 27599-7264; Curriculum for Bioinformatics and Computational Biology, University of North Carolina, Chapel Hill, 120 Mason Farm Road, Chapel Hill, NC 27599-7264; Lineberger Comprehensive Cancer Center, University of North Carolina, Chapel Hill, 120 Mason Farm Road, Chapel Hill, NC 27599-7264

**Keywords:** cell cycle, live-cell imaging, single-cell dynamics, computational modeling, scientific visualization

## Abstract

**SUMMARY:** The cell cycle is driven by precise temporal coordination among many molecular activities. To understand and explore this process, we developed the Cell Cycle Browser (CCB), an interactive web interface based on real-time reporter data collected in proliferating human cells. This tool facilitates visualizing, simulating, and predicting the outcomes of perturbing cell cycle parameters. Time-series traces from individual cells can be combined to build a multi-layered timeline of molecular activities. Users can simulate the cell cycle using computational models that capture the dynamics of molecular activities and phase transitions. By adjusting individual expression levels and strengths of molecular relationships, users can predict effects on the cell cycle. Virtual assays, such as growth curves and flow cytometry, provide familiar outputs to compare cell cycle behaviors for data and simulations. The CCB serves to unify our understanding of cell cycle dynamics and provides a platform for generating hypotheses through virtual experiments.

**HIGHLIGHTS:**

- Users can stack and align single-cell traces for different molecular reporters
- Computational models with adjustable parameters simulate cell cycle progression
- Virtual growth curves and flow cytometry assays predict cell cycle behaviors

## MAIN TEXT

The cell-division cycle—the process by which cells replicate their DNA to form two daughter cells—is central to nearly every facet of the life sciences including disease research and industrial biotechnology. A long-term challenge for the field is to determine precisely how the cell cycle is coordinated in time and how that coordination changes under different environmental conditions. A growing number of recent studies have monitored various molecular entities in single cells over the course of the cell cycle including the activities of cyclin-dependent kinases (Spencer et al., 2013), stress response factors (Loewer et al., 2010), DNA damage repair foci (Arora et al., 2017), and DNA replication (Barr et al., 2017, Chao et al., 2017). A high-resolution timeline that accounts for the interrelationships among cell cycle entities would be a transformative tool for many basic and applied fields of research. Such a map would predict how future changes—such as a newly discovered mutation, environmental exposure, or drug—are likely to affect cell proliferation and genome stability. Here, we describe a web-based tool, the Cell Cycle Browser (CCB), for visualizing, simulating, and analyzing the timeline of molecular events during cell cycle progression (https://cellcycle.renci.org). This article provides a high-level overview of the CCB. A more thorough tutorial of the tool is available at https://github.com/RENCI/CellCycleBrowser/wiki/tutorial.

### CCB Workspace

The information displayed by the CCB is organized into *workspaces.* A workspace consists of a collection of *datasets*—experimentally measured, single-cell traces of molecular activities—and cell cycle *models*—files describing the kinetics relationships between molecular activities and the rates of cell cycle progression. Cell cycle models can be simulated and compared to experimental data. In the data selection section of the CCB graphical user interface (GUI) (**Figure 1-1**), the user can select from a list of available workspaces. After selecting a workspace, the datasets and first model included in that workspace are automatically loaded. The user can add additional datasets, remove any currently loaded datasets, or select a different model.

**Figure 1.**
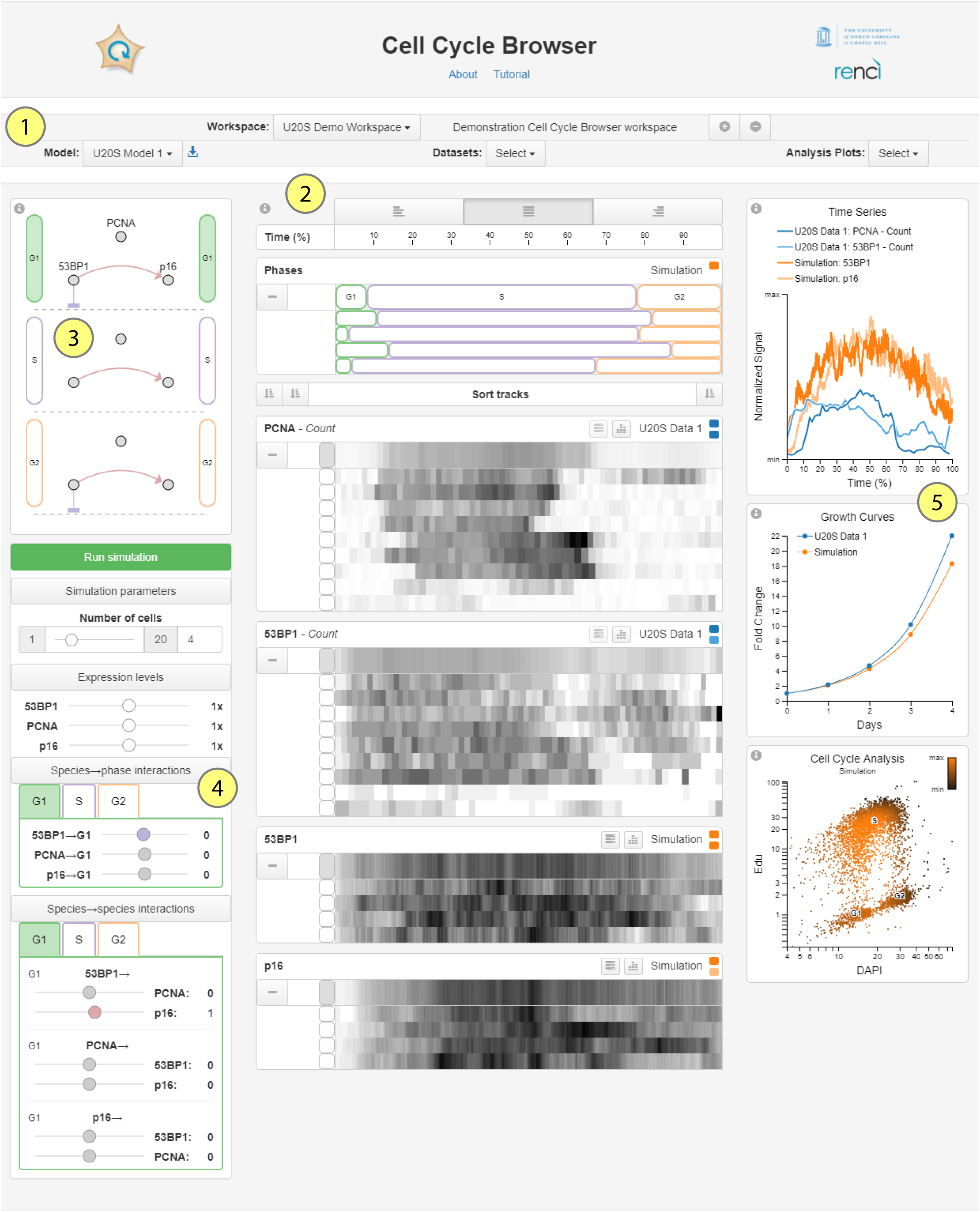
Cell Cycle Browser user interface. 1. Data selection section. 2. Data browser section. 3. Map visualization. 4. Model parameter controls. 5. Analysis plots.

Each dataset contains a time-series of measurements for any number of molecular species captured across multiple individual cells. Each single-cell time-series, or trace, is derived from the analysis of segmented images of proliferating human cells expressing a fluorescent reporter. Therefore, each species may contain multiple features such as reporter intensity, number of foci, nuclear area, or the spatial position of the cell. The available features for each dataset can be selected from the dropdown menu, and are displayed in the Data Browser. The user can select a single model from the dropdown model list, or choose “None” if interested in viewing only the experimental datasets. Finally, the user can select which analysis plots to display including a time-series plot of experimental and simulated data; a growth curve plot for each dataset/simulation; or a simulated analytical flow cytometry plot of DNA synthesis vs. DNA content (FACS) for the current simulation or any datasets that contain phase information.

### Data Browser

One of the primary uses of the CCB is that it provides a common time axis to visualize multiple time-series traces from individual cells representing diverse molecular activities. The center section of the CCB provides a heat-map representation of single-cell traces from the current experimental data and/or simulated output (**Figure 1-2**). To capture a single cell cycle, each cell’s trace is excised from the telophase event that generated that cell until its own telophase (Davis and Purvis, 2014). A separate *track* is drawn for each selected image feature across all molecular species in a particular data *source*. A data source refers to an experimental dataset, which may contain multiple reporters and image features, or the current model simulation. Each track consists of a collapsible area showing each individual cell trace with value mapped to color (white: low, black: high). Color scaling, which performs unit-normalization on the time-series values, can be performed per track, or per trace. An average trace for each track is displayed above the single-cell traces. Average traces are generated by scaling the length of each individual trace to match the average length of all traces per track and then performing nearest-neighbor sampling for each sample point in the average. Traces can be aligned to the beginning (first telophase), the end (second telophase), or individually stretched to align each trace to the beginning and end of the cell cycle.

The heat-map visualization facilitates differentiation of individual traces while still supporting analysis of different temporal patterns (Correll et al., 2012, Albers et al., 2014). In addition to molecular activities, the CCB draws tracks showing the locations of individual cell cycle phases (G1, S, G2-M) both for the simulated data as well as for any datasets that contain phase information. Tracks can be sorted by type (i.e., data or phase), species, image feature, or data source. The user can also arrange tracks by dragging and dropping them individually. Buttons to the left of each trace control whether or not to show that trace in the time-series analysis plot, and colored icons in the track header indicate the color for that data source/track in any displayed *Analysis Plots* (described below).

### Analysis Plots

Three analysis plots are available to provide additional information about the experimental and simulated data (**Figure 1-5**). A time series plot shows each individual or average cell trace—selected using buttons in the data browser tracks—as a line plot. Two or more time series plots from the same source have the same base color, with different shades of that color for each track from the data source. These colors are also indicated by the color icons in each track; the data source color is used for the other analysis plots. A growth curve plot shows fold change growth per day for each data source, based on the average length of the traces for that track. Flow cytometry plots (DNA synthesis on the y axis and DNA content on the x-axis) are simulated for each data source containing phase information. Each data point represents a simulated cell, with phase generated probabilistically based on the average phase lengths for that data source (Toettcher et al., 2009). Each cell is colored based on the local density of neighboring cells.

### Model Visualization

The left section of the CCB contains a network visualization of the cell cycle model (**Figure 1-3**) as well as controls for adjusting the model and simulation parameters (**Figure 1-4**). In a model, each molecular activity is referred to as a *species*—a pool of entities that are assumed to be indistinguishable, located in the same cellular compartment, and participate in the same molecular reactions (Hucka et al., 2003). Each cell cycle model specifies: (1) the abundance of each species (e.g., activity of Cdk2; number of 53BP1 foci); (2) the influence of each species on each phase transition (e.g., influence of Cdk2 activity on the G1/S transition; inhibitory effect of 53BP1 damage foci on G2/M transition); and (3) the influence of each species on every other species for each phase (e.g., an increase in 53BP1 foci during G2 induces expression of p21).

In the network diagram, each species is represented as a circular node whose size is proportional to the species’ abundance (specifically, the fold change compared to the initial value specified in the model). A separate species diagram is drawn for each phase since interactions may vary throughout the cell cycle. Links between two species or between a species and a particular phase transition indicate the interaction between those entities, color-mapped from blue (inhibiting) through grey (no influence) to red (activating), with magnitude mapped to width. Selecting a particular phase in the visualization activates the control tabs for that phase in the interaction panels, described below.

### Model Simulation and Perturbation

A set of computational models describing cell cycle progression have been developed for different cell types and molecular activities. These models can be simulated under the default conditions or after adjusting model parameters. Simulation parameters can be adjusted using sliders that control: a) number of cells to simulate, b) species expression levels, c) species to phase interaction parameters, and d) species-to-species interaction parameters per phase. Interaction slider handles are colored to match the links in the network diagram visualization.

When the simulation button is pressed, the StochPy library (Maarleveld et al., 2013) is used to perform cell cycle simulations on the server via the stochastic simulation algorithm (SSA) (Gillespie, 1977, Gillespie, 2007). A customized StochPy simulation engine terminates the simulation when the specified last phase of the cell cycle (e.g., M-phase) is completed in order to simulate exactly one complete cell cycle per simulation. Because some simulation parameters can cause extremely slow cell cycle simulation (e.g., high expression of p21), a red cancel button is available so users can cancel the run if the simulation is taking an excessively long time.

In summary, the Cell Cycle Browser (CCB) (https://cellcycle.renci.org) is a web-based tool for exploring, simulating, and perturbing the human cell cycle via an interactive graphical user interface (GUI). It was developed in the spirit of the UCSC Genome Browser (https://genome.ucsc.edu), which provides a graphical view of genome sequence data along the physical coordinates of the chromosomes. Unlike the Genome Browser, however, the primary axis of the CCB is time. In its most basic functionality, the CCB displays time series data representing various molecular activities over the course of a single cell cycle. Single-cell traces from time-lapse fluorescent microscopy experiments – representing different biological activities from multiple cell lines – may be aligned in time to examine the temporal sequence of cell cycle events and explore new relationships among cell cycle events. In addition, the CCB enables a user to simulate a mechanistic model of cell cycle progression and align the model output with experimental data in time to better understand and predict cell cycle behaviors.

Future improvements to the CCB will include: (1) the ability to create customized, password-protected workspaces for individual users that allows them to create private workspaces for intermediate work, which they can choose to make public later; (2) data from additional organisms; (3) a portal for submitting raw data to be available for visualization as tracks; (4) automated imputation of phase transition locations based on single-cell traces; and (5) the ability to load, simulate, and compare multiple computational models.

## STAR*METHODS

KEY RESOURCES TABLE

**KEY RESOURCES TABLE**
| RESOURCE | SOURCE | IDENTIFIER |
| --- | --- | --- |
| Sample Data |  |  |
| U2OS cell dataset | Chao et al., 2017 | https://cellcycle.renci.org/cell_data_meta/PCNA_53BP1.csv/ |
| Software and Algorithms |  |  |
| CCB tool software<br>source code repo | This paper | https://github.com/RENCI/CellCycleBrowser/ |
| Docker | https://www.docker.com/ | v1.7.1 |
| Django Python web<br>framework | https://www.djangoproject.com/ | v1.10 |
| Python | https://www.python.org/ | v2.7.11 |
| PostgreSQL database | https://www.postgresql.org/ | v2.6.1 |
| RabbitMQ message<br>broker | https://www.rabbitmq.com/ | v1.4.9 |
| Celery distributed task<br>queue | http://www.celeryproject.org/ | v3.1.23 |
| StochPy stochastic<br>modeling in Python | http://stochpy.sourceforge.net/ | v2.3 |
| libSBML Python library | http://sbml.org/Software/libSBML/docs/python-api/ | v5.15.0 |
| SimpleSBML library | http://www.biorxiv.org/content/early/2015/10/30/030312 | v1.1 |
| Bootstrap | http://getbootstrap.com/ | v3.3.6 |
| jQuery | https://jquery.com/ | v1.12.4 |
| d3 | https://d3js.org/ | v4.1.0 |
| d3-contour<br>d3-scale-chromatic |  |  |
| flux | https://github.com/facebook/flux | v2.1.1 |
| keymirror | https://github.com/STRML/keyMirror | v0.1.1 |
| object-assign | https://github.com/sindresorhus/object-assign | v4.1.0 |
| react<br>react-dom | https://facebook.github.io/react/ | v 15.2.0 |
| science | https://github.com/jasondavies/science.js/ | v1.9.3 |
| seedrandom | https://github.com/davidbau/seedrandom | v2.4.3 |
| Other |  |  |
| Website for CCB portal | This paper | https://cellcycle.renci.org |
| Tutorial for CCB tool | This paper | https://github.com/RENCI/CellCycleBrowser/wiki/Tutorial |
| CCB installation guide<br>for developers | This paper | https://github.com/RENCI/CellCycleBrowser/blob/master/README.md |

## CONTACT FOR DATA SHARING

Please contact Jeremy Purvis at with requests and inquiries.

## METHOD DETAILS

### Server handling of data and model simulation

JSON was used as the data interchange format to pass AJAX updates between the client and the server. **Figure S1** shows a pipeline of detailed data flow to illustrate how data flows between the client and the server via AJAX when the page initially loads and at different requests from user interactions with the browser.

When the main page initially loads, an AJAX request is sent from the client and the server responds with an array of key-value pairs for all workspaces (collections of datasets and models) in JSON format. Key-value pairs for each workspace include name and description of the workspace along with an array of key-value pairs for all datasets and models included in the workspace. The key-value pairs for each dataset or model include name, description, and the name of the file stored on the server. The client also requests from the server via AJAX a list of all datasets and a list of all models as needed to populate the user interface.

Each experimental dataset is stored on the server as a series of time-stamped values of features for one or more species in CSV format, with metadata included in a header section. Each model input file is stored on the server in SBML (Systems Biology Markup Language) format. When a dataset is selected, the client sends AJAX requests to the server to retrieve time-stamped data for the selected dataset for visualization. When a model is selected, a corresponding AJAX request is sent and the server extracts initial parameter data from the model and sends the data in JSON format to the client for display in the model control panel and to construct and display the map visualization. These model initial parameter data include species expression levels, species-phase interactions, and species-species interactions per phase; all can be adjusted from the user interface before triggering a model simulation run.

When a model simulation run is triggered from the user interface, the client sends the simulation request to the server via AJAX, including all input parameters and the number of cells/trajectories to simulate. The server performs the stochastic simulation algorithm (SSA) as an asynchronous task in a separate worker thread on the web server without blocking the main web thread. Utilizing multithreading and asynchronous model simulation runs makes the CCB web tool responsive, scalable, and performant even with large numbers of simultaneous web requests for model simulation runs. When the simulation is complete, the server sends the simulation result to the client as an AJAX response so that the simulation result can be visualized and analyzed to compare with experimental datasets.

### Model Specification

Models were specified using the Systems Biology Markup Language (SBML) (Hucka et al., 2003). We used SimpleSBML (Cannistra et al., 2015) to write SBML models using the following user-specified parameters:

1. The name of the output file (e.g., “model.xml”).
2. *k* and *λ* values for each cell cycle phase using the Erlang model in Chao et al. (Chao et al., 2017).
3. A list of *n* species names, S.
4. A list of *n* initial conditions, X0. Values must be >= 0.
5. An *n*-by-*m* array of species production rates in each phase, B, where *m* is the number of phases. Values must be >= 0.
6. An *n*-by-*m* array of species degradation rates in each phase, A. Values must be >= 0.
7. An *n*-by-*m* array of species-phase interaction strengths, P. These values indicate how each species increases or decreases the rate of progression through a particular phase. Values can be positive, negative, or zero. A value of 0 does not change the rate of progress. Values less than 0 slow progress and values greater than 0 increase the rate of progress.
8. An *n*-by-*n*-by-*m* array of species-species interactions for each phase, Z. Each value Z_*ijk*_ represents the strength of the influence of species *i* on species *j* during phase *k*. Values can be positive, negative, or zero. Positive values indicate promoting interactions; negative values indicate negative interactions. A value of 0 indicates no interaction.

Using these values, SimpleSBML generates an XML file using the following steps:

1. Generate an “invisible” species for each phase (e.g., “G1”) and subphase (e.g., “G1_01”, “G1_02”,…). Phase and subphase values can be either zero or one. Only one subphase can be one; all others must be zero.
2. Generate an assignment rule so that each phase is the sum of all of its subphases. In this way, each phase/suphase variable is an indicator of the current phase/subphase.
3. Generate reactions to transition between each phase (e.g., consuming G1_01 and producing G2_02 at rate *λ*_G1_)
4. Generate each species in list S and set its initial condition to values in X0.
5. For each species, S_i_, generate a production reaction: null -> S_i_. Set the rate law to: Bi1*(G1) + Bi2*(S) + Bi3*(G2), where Bi1 refers to the production term for the ith species for the G1 phase (the first column in the array, B). Since G1, S, and G2 are indicator variables, this ensures that Si is produced at rate Bi1 only when the cell is in G1.
6. For each species, Si, generate a degradation reaction: Si -> null. Set the rate law to: Ai1*(G1)*Si + Ai2*(S)*Si + Ai3*(G2)*Si, where Ai1 refers to the degradation term for the ith species for the G1 phase (the first column in the array, A). Since G1, S, and G2 are indicator variables, this ensures that Si is degraded at rate Ai1 only when the cell is in G1.
7. For each element Pij in array P, add Pij to each of the reactions for phase transition j. For example, if Pij represents the effect of 53BP1 on S phase, then all of the S phase transition reactions (e.g., S1 -> S2) should have rate law: *λ*_S_ * power(1+S1, Pij). Here, *λ* is the rate already specified in the first 6 arguments that define basal transition rates. Pij will speed up or slow down this rate.
8. For each non-zero element Zijk, generate a reaction. If Zijk > 0, generate a production reaction: null -> Sj with rate law: Zijk * Si. If Zijk < 0, generate a degradation reaction: Sj -> null with rate law: Zijk * Si * Sj.

### Client/GUI architecture

The client was built with JavaScript and HTML using the React library for creating custom GUI components, Bootstrap for GUI styling, and D3 for creating the different SVG-based visualization elements. The Flux architecture is used to ensure consistent dataflow.

### Model parameters

There are two types of model parameters that can be adjusted by the user via sliders: species expression levels and interactions between species and phases, and between species. The expression level for each species in the initial model is the basal level for that protein. As such, these may differ between species. The slider controls for the expression level impart a fold-change on this initial expression level, with each slider initially at a value of 1x. Sliding to the right imparts a fold increase, up to 32x, and to the left a fold decrease, ending in 0x, which in effect removes the species. The interaction sliders describe the exponent in the rate equations for each slider. These sliders range from -10 to 10, where more negative numbers apply an increasingly inhibiting interaction, and more positive numbers apply an increasingly promoting interaction.

### Averaging traces

To calculate the average trace for each track, the average time steps across all traces in that track is calculated and used to generate sample points up to the average trace length. Each trace is then stretched temporally to match the average trace length, and nearest-neighbor interpolation is used to average across all traces for each sample point.

### Growth curve calculation

Growth curves for each data source (experimental dataset or simulation) are shown in the Growth Curve Analysis Plot. The growth curves are calculated from the average trace length in that data source with the formula *y* = 2^*t*/*l*^ where *y* is the fold change in cell number, *t* is the independent variable of time on the x-axis of the growth curve, and *l* is the average duration of the cell cycle for a group of single-cell traces.

### Cell cycle analysis plots

A virtual flow cytometry cell cycle analysis was performed for each data source containing phase information and displayed in the Cell Cycle Analysis Plot. Separate scatter plots of the steady-state of a large number of cells are generated by resampling a kernel-density estimate of flow cytometry from an experimental dataset, based on the phase lengths in the given data source. The phase probability distributions for each data source are calculated based on the appropriate formulas in (Toettcher et al., 2009). First, we calculate τ_phase_, the normalize length of each phase, by dividing the average length for that phase by the average cell cycle length for that data source. The proportion of cells, P_phase_, in each phase are then:

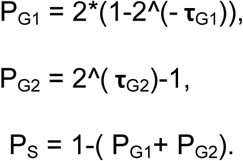

These probabilities are used to generate the phase for each simulated cell, and then resampling of the kernel density estimate for that phase is used to generate an (x, y) point in the resulting scatterplot.

## AUTHOR CONTRIBUTIONS

D.B. and H.Y. developed the user interface and server-side software, contributed to the conceptual design of the CCB, and wrote portions of the manuscript. G.G. conceptualized the virtual experiments and contributed live-cell data. K.K. and H.X.C contributed live-cell data, performed image analysis, and standardized data formatting. R.A.H. developed the logo and contributed to visualization and image analysis. J.K. developed SBML models to simulate cell cycle progression. S.W. performed time-lapse imaging and contributed live-cell data. J.G.C. proposed virtual experiments and improvements to the user interface and contributed live-cell data. J.E.P. conceived of the CCB and wrote portions of the manuscript.

## ACKNOWLEDGEMENTS

We thank Sabrina Spencer and Mansi Arora for contributing published live-cell data to the browser. This work was supported by a Medical Research Grant from the W.M. Keck Foundation (J.E.P. and J.G.C.), NIH pre-doctoral fellowships F30-CA213876 (H.X.C.) and F31-HL134336 (R.A.H.)

---

**Figure S1.**
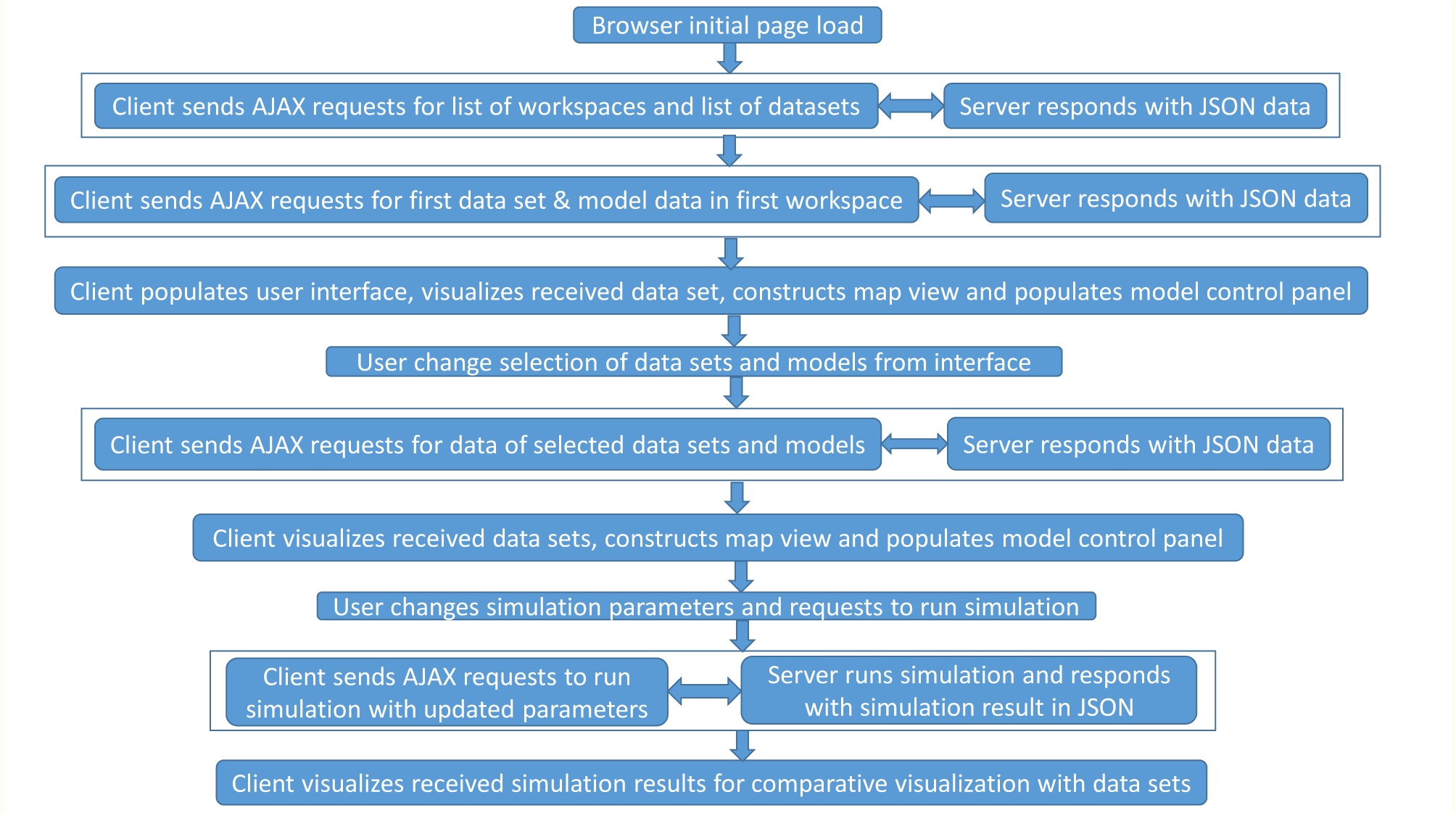
Pipeline for data flow, Related to Figure 1.

